# *Bacteroides thetaiotaomicron* metabolic activity decreases with polysaccharide molecular weight

**DOI:** 10.1101/2023.09.21.558885

**Authors:** Jeremy P. H. Wong, Noémie Chillier, Michaela Fischer-Stettler, Samuel C. Zeeman, Tom J. Battin, Alexandre Persat

**Affiliations:** Institute of Bioengineering and Global Health Institute, School of Life Sciences, École Polytechnique Fédérale de Lausanne, Lausanne, Switzerland; School of Architecture, Civil and Environmental Engineering, École Polytechnique Fédérale de Lausanne, Sion, Switzerland; Department of Biology, ETH Zürich, 8092 Zürich, Switzerland

## Abstract

The human colon hosts hundreds of commensal bacterial species, many of which ferment complex dietary carbohydrates. To transform these fibers into metabolically-accessible compounds, microbes often express series of dedicated enzymes homologous to the starch utilization system (sus) encoded in polysaccharide utilization loci (PULs). The genome of *Bacteroides thetaiotaomicron (Bt),* a common member of the human gut microbiota, encodes nearly 100 PULs, conferring a strong metabolic versatility. While the structures and functions of individual enzymes within the PULs have been investigated, little is known about how polysaccharide complexity impacts the function of sus-like systems. We here show that the activity of sus-like systems depends on polysaccharide size, ultimately impacting bacterial growth. We demonstrate the effect of size-dependent metabolism in the context of dextran metabolism driven by the specific utilization system PUL48. We find that as molecular weight of dextran increases, *Bt* growth rate decreases and lag time increases. At the enzymatic level, the dextranase BT3087 is the main glycosylhydrolase for dextran utilization and that BT3087 and BT3088 contribute to *Bt* dextran metabolism in a size-dependent manner. Finally, we show that the polysaccharide size-dependent metabolism of *Bt* impacts its metabolic output in a way that modulates the composition of a producer-consumer community it forms with *Bacteroides fragilis*. Altogether, our results expose an overlooked aspect of *Bt* metabolism which can impact the composition and diversity of microbiota.

## Introduction

Polysaccharides are complex, polymeric molecules composed of long chains of monosaccharide subunits linked together through glycosidic linkages^1^. These molecules are essential to life^2^ and are commonly found in our daily diets^3^. Their chemical structures are variable due to differences in monosaccharide compositions and glycosidic linkage patterns^4^. As a result, the degradation of these molecules requires diverse sets of enzymes that generally target specific polysaccharides. While humans lack the ability to degrade many polysaccharides, their intestinal microbiota contains bacterial commensals that are versatile polysaccharide utilizers. In fact, microbiota species are selected for the ability to forage more or less complex carbohydrates and glycans^5^. Thus, for many gut microbes, the ability to persist in the gut correlates with their ability to utilize dietary fibers and other carbohydrate-based nutrients^6^. Therefore, understanding the mechanisms of polysaccharide utilization by gut microbes may provide an understanding to the forces driving the composition of the gut microbiota^5^.

*Bacteroides spp.* are among the most abundant and well-studied glycan-foraging commensals found in the human bowel^7^. *Bacteroides* can utilize complex carbohydrates through the expression of series of protein systems encoded in polysaccharide utilization loci (PULs)^8^. PULs encode for proteins which bind, cleave and import complex polysaccharides into simpler, metabolically-accessible sugars^9–13^. *Bacteroides thetaiotaomicron (Bt)* can utilize many dietary polysaccharides including starch^14^, arabinan^15^, levan^16^, inulin^17^, rhamnogalacturonan II^18^, complex dietary N-glycans^19^, and even glycans derived from its mammalian hosts such as o-linked glycans from mucins and HMO (human milk oligosaccharides)^20^.

The starch utilization system (Sus) has served as model system for PUL studies^14^. In *Bt*, 8 genes encode Sus (*susRABCDEFG*). SusD, SusE and SusF are outer membrane lipoproteins which are responsible for the initial binding of oligosaccharides to the cell surface^14,21^. SusG is an outer membrane-associated α-amylase which hydrolyzes the starch into maltooligosaccharides^22^. The resulting maltooligosaccharides enter the periplasm thanks to the TonB-dependent transporter SusC^23^. These sugars are further cleaved by two periplasmic glycosylhydrolases (GHs) SusA (neopullulanase) and SusB (α-glucosidase)^24,25^. The inner membrane-spanning sensor-regulator protein SusR regulates the expression Sus genes in response to periplasmic levels of maltose^26^. Other PULs are Sus homologs encoding the nutrient binding and importer proteins SusD and SusC, lipoproteins of similar structures to SusE and SusF, along with a series of diverse GHs dedicated to a wide range of different complex carbohydrates, which are regulated by a SusR-like protein^27^. The genome of *Bt* encode 96 different PULs^28^, allowing them to grow on a broad range of dietary or host-derived glycans^9,10,12,19^.

Although different PULs are dedicated to the degradation of specific polysaccharides with unique glycosidic linkages and monosaccharide compositions, a specific polysaccharide may be found with different complexities such as molecular weight (MW) and degree of branching^4^, as in starch produced by different plants^29^. Sus-like systems are composed of multiple surface-exposed components that need to coordinate functions to properly cleave and import degraded oligosaccharides. These must therefore handle polymers with the vastly differing physical attributes which may impact their processability. However, little is known about how MW impacts the efficacy of PULs in producing accessible glycans that act as accessible substrate for growth.

In a previous work^30^, we have identified dextran as a polysaccharide readily utilized by *Bt*. Like starch, dextran is a polyglucan consisting of 2 unique glycosidic linkages: an α-1,6 linked backbone along with α-1,3 branches^31^. Dextran is a common additive in the food industry^32^. Unlike starch, dextran polymers are highly soluble in water. Also, due to their extensive commercial and clinical applications, there exists many purification protocols which produce dextran polymers of defined MW^32^. Using *Bt* as model commensal and dextran as model dietary fiber, we investigated the impact of polysaccharide MW on the efficacy of dextran utilization by *Bt*. We found that *Bt* growth rate decreases as dextran MW increased. By studying dextran metabolism with *Bt* mutants containing knockouts of PUL48 genes, we identified protein targets within PUL48 whose importance in dextran utilization is MW-dependent, thereby suggesting a mechanism of size-dependent metabolism. We ultimately demonstrate that size-dependent metabolism has an impact on the composition of a model microbiota community.

## Results

### Bt uses PUL48 to degrade dextran

*Bt* exposed to dextran as sole carbon source upregulates proteins belonging to the predicted PUL48 and the α-glucosidase BT4581^30^ (Fig. 1A). Consistent with this, a functional genomic screen identified genes from PUL48 to be essential for growth in dextran^33^. PUL48 comprises of a SusD-like surface associated nutrient binding protein (BT3089), a SusC-like TonB-dependent importer protein (BT3090), a SusR-like sensor-regulator protein (BT3091) and 2 GHs: α-glucosidase II (BT3086) and dextranase (BT3087) (Fig. 1A). To study the functions of individual PUL48 proteins in *Bt* dextran metabolism, we generated knockout mutants of PUL48 genes (Table 1). We first characterized the essentiality of these genes for growth in dextran. We grew individual mutants in *Bacteroides* Minimal Medium (BMM) supplemented with dextran (MW = 35 kDa) over 3 days (Fig. 1B). We observed that *Bt* mutants lacking the α-glucosidase II BT3086 and nutrient binding protein BT3088 (SusE) resulted in marginal growth defects in dextran. Mutants lacking the α-glucosidase BT4581 showed delayed growth. Mutants in nutrient binding protein gene BT3089 (SusD) and importer protein BT3090 (SusC) showed severely impaired growth in dextran in comparison to WT *Bt*. Finally, *Bt* mutants in the regulatory protein BT3091 (SusR) and the dextranase BT3087 were entirely unable to grow in dextran (Fig. 1B). We examined the growth profiles of *Bt* mutants lacking several GHs: *Δ86,81* (double deletion of BT3086 and BT4581)*, Δ87,81* (double deletion of BT3087 and BT4581)*, Δ86,87* (double deletion of BT3086 and BT3087) *and Δ86,87,81* (triple deletion of BT3086, BT3087 and BT4581 also suggest that cell growth is eliminated only in the case of the loss of dextranase BT3087 (Fig. S1A). These results suggests that the proteins BT3086, BT3088 and BT4581 are auxiliary proteins which promote but are not essential to dextran fermentation. The proteins BT3089 (SusD) and BT3090 (SusC) are important for the proper growth of *Bt* in dextran. Their absence likely leads to a decrease in nutrient availability to *Bt* through impaired oligosaccharide binding and import.

**Figure 1:**
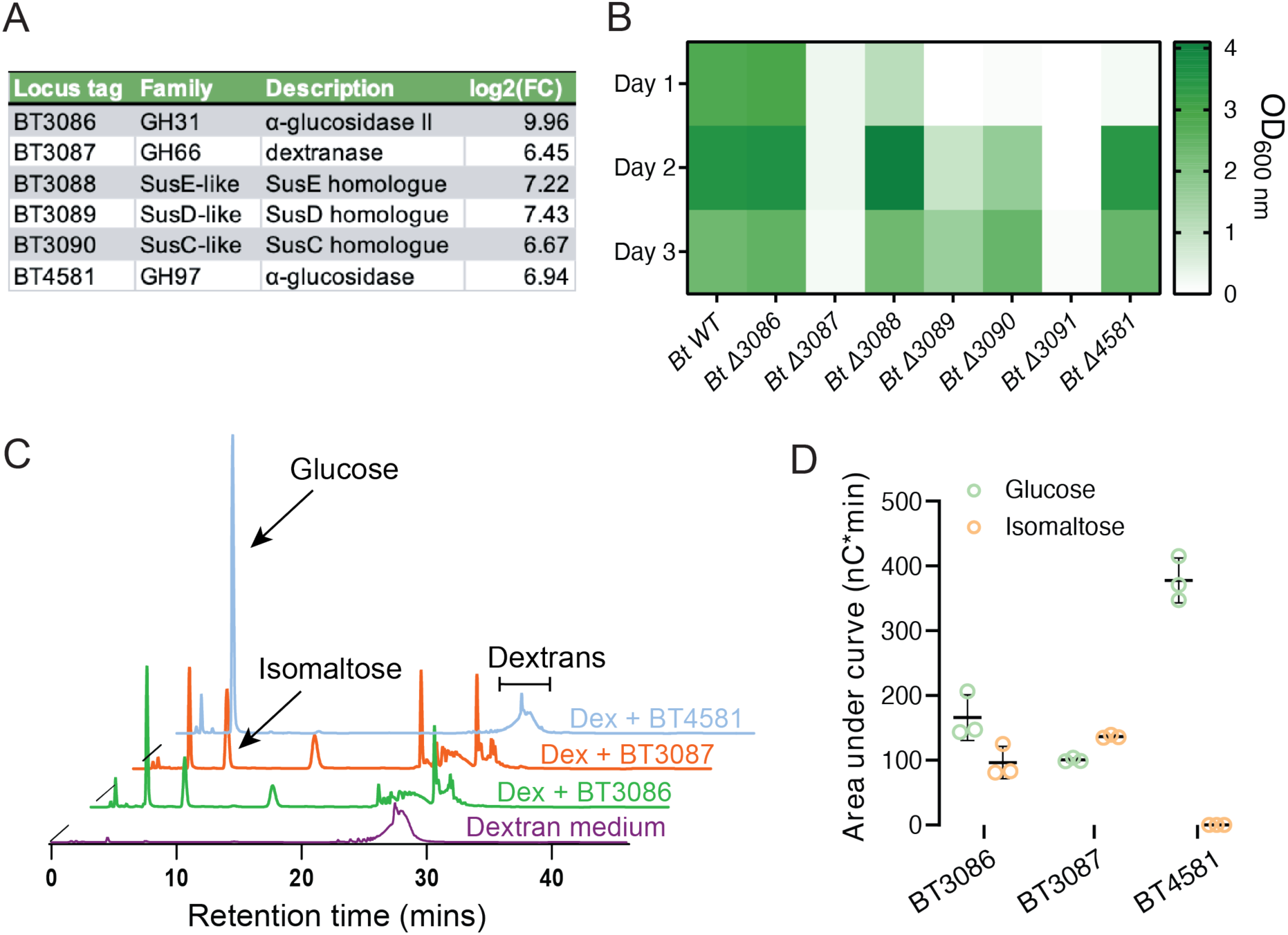
The roles of PUL48 proteins in the mechanism of dextran metabolism. (A) List of proteins (green dots) which are significantly upregulated in dextran along with their functions. These proteins include 6 proteins belonging to the predicted PUL48, along with a sole GH BT4581. (B) Monoculture growths of *Bt* WT and various PUL48 KO mutants in dextran. We measured the optical density (OD) of individual cultures everyday over the course of 3 days. (C) HPAEC-PAD analysis of dextran medium treated overnight with recombinantly expressed GHs: BT3086, BT3087 or BT4581. Violet = no treatment, Green = treatment with BT3086, orange = treatment with BT3087 and blue = treatment with BT4581. (D) Area under the curve analysis of the glucose and isomaltose peaks of dextran treated with BT3086, BT3087 or BT4581. BT3086 and BT3087 form both glucose and isomaltose as products, whereas BT4581 forms solely glucose. (D-E) HPAEC-PAD experiments were run in triplicates, we display one representative chromatogram.

**Table 1.** List of strains and plasmids generated for this study.

| Reagent type | Background | Designation | Source | Description |
| --- | --- | --- | --- | --- |
| Strain | *B. thetaiotaomicron*<br>VPI-5482 | *Bt* WT | <sup>45</sup> | *Bt* wildtype |
| Strain | *B. thetaiotaomicron*<br>VPI-5482 | *Bt* WT - sfGFP | <sup>45</sup> | *Bt* wildtype<br>constitutively<br>expressing sfGFP |
| Strain | *B. thetaiotaomicron*<br>VPI-5482 | *Bt* WT - mCherry | <sup>45</sup> | *Bt* wildtype<br>constitutively<br>expressing mCherry |
| Strain | *B. thetaiotaomicron*<br>VPI-5482 | *Bt* ΔBT3086 | This<br>study | Deletion of α-<br>glucosidase II<br>BT3086 |
| Strain | *B. thetaiotaomicron*<br>VPI-5482 | *Bt* ΔBT3087 | This<br>study | Deletion of<br>dextranase BT3087 |
| Strain | *B. thetaiotaomicron*<br>VPI-5482 | *Bt* ΔBT3088 | This<br>study | Deletion of SusE like<br>BT3088 |
| Strain | *B. thetaiotaomicron*<br>VPI-5482 | *Bt* ΔBT3089 | This<br>study | Deletion of SusD like<br>BT3089 |
| Strain | *B. thetaiotaomicron*<br>VPI-5482 | *Bt* ΔBT3090 | This<br>study | Deletion of SusC like<br>BT3089 |
| Strain | *B. thetaiotaomicron*<br>VPI-5482 | *Bt* ΔBT3091 | This<br>study | Deletion of SusR like<br>BT3089 |
| Strain | *B. thetaiotaomicron*<br>VPI-5482 | *Bt* ΔBT4581 | This<br>study | Deletion of α-<br>glucosidase BT4581 |
| Strain | *B. thetaiotaomicron*<br>VPI-5482 | *Bt* ΔBT3086 sfGFP | This<br>study | *Bt* ΔBT3086<br>constitutively<br>expressing sfGFP |
| Strain | *B. thetaiotaomicron*<br>VPI-5482 | *Bt* ΔBT3087 sfGFP | This<br>study | *Bt* ΔBT3087<br>constitutively<br>expressing sfGFP |
| Strain | *B. thetaiotaomicron*<br>VPI-5482 | *Bt* ΔBT3088 sfGFP | This<br>study | *Bt* ΔBT3088<br>constitutively<br>expressing sfGFP |
| Strain | *B. thetaiotaomicron*<br>VPI-5482 | *Bt ΔBT3089* sfGFP | This study | *Bt ΔBT3089*<br>constitutively<br>expressing sfGFP |
| Strain | *B. thetaiotaomicron*<br>VPI-5482 | *Bt ΔBT3090* sfGFP | This study | *Bt ΔBT3090*<br>constitutively<br>expressing sfGFP |
| Strain | *B. thetaiotaomicron*<br>VPI-5482 | *Bt ΔBT3091* sfGFP | This study | *Bt ΔBT3091*<br>constitutively<br>expressing sfGFP |
| Strain | *B. thetaiotaomicron*<br>VPI-5482 | *Bt ΔBT4581* sfGFP | This study | *Bt ΔBT4581*<br>constitutively<br>expressing sfGFP |
| Strain | *B. thetaiotaomicron*<br>VPI-5482 | *Bt ΔBT3088*<br>mCherry | This study | *Bt ΔBT3088*<br>constitutively<br>expressing mCherry |
| Strain | *B. thetaiotaomicron*<br>VPI-5482 | *Bt ΔBT3089*<br>mCherry | This study | *Bt ΔBT3089*<br>constitutively<br>expressing mCherry |
| Strain | *B. thetaiotaomicron*<br>VPI-5482 | *Bt ΔBT3090*<br>mCherry | This study | *Bt ΔBT3090*<br>constitutively<br>expressing mCherry |
| Strain | *B. thetaiotaomicron*<br>VPI-5482 | *Bt Δ86,87* | This study | Deletion of BT3086<br>and BT3087 |
| Strain | *B. thetaiotaomicron*<br>VPI-5482 | *Bt Δ86,81* | This study | Deletion of BT3086<br>and BT4581 |
| Strain | *B. thetaiotaomicron*<br>VPI-5482 | *Bt Δ87,81* | This study | Deletion of BT3087<br>and BT4581 |
| Strain | *B. thetaiotaomicron*<br>VPI-5482 | *Bt Δ86,87,81* | This study | Deletion of BT3086,<br>BT3087 and BT4581 |
| Strain | *B. thetaiotaomicron*<br>VPI-5482 | *Bt Δ86,87* sfGFP | This study | *Bt Δ86,87*<br>constitutively<br>expressing sfGFP |
| Strain | *B. thetaiotaomicron*<br>VPI-5482 | *Bt Δ86,81* sfGFP | This study | *Bt Δ86,81*<br>constitutively<br>expressing sfGFP |
| Strain | *B. thetaiotaomicron*<br>VPI-5482 | *Bt* Δ87,81 sfGFP | This study | *Bt* Δ87,81 constitutively expressing sfGFP |
| Strain | *B. thetaiotaomicron*<br>VPI-5482 | *Bt* Δ86,87,81 sfGFP | This study | *Bt* Δ86,87,81 constitutively expressing sfGFP |
| Strain | *B. fragilis* NCTC-9343 | *Bf* mCherry | <sup>45</sup> | *Bf* constitutively expressing mCherry |
| Plasmid | - | Λpir pWW3452 | <sup>45</sup> | Plasmid for insertion of sfGFP into *Bt* |
| Plasmid | - | Λpir pWW3515 | <sup>45</sup> | Plasmid for insertion of mCherry into *Bt* |
| Plasmid | pLGB13 | pLGB13 | <sup>44</sup> | Plasmid backbone for *Bt* knockouts |
| Plasmid | pLGB13 | pAB001 | This study | Plasmid for the deletion of *BT3086* |
| Plasmid | pLGB13 | pAB002 | This study | Plasmid for the deletion of *BT3087* |
| Plasmid | pLGB13 | pAB003 | This study | Plasmid for the deletion of *BT3088* |
| Plasmid | pLGB13 | pAB004 | This study | Plasmid for the deletion of *BT3089* |
| Plasmid | pLGB13 | pAB005 | This study | Plasmid for the deletion of *BT3090* |
| Plasmid | pLGB13 | pAB006 | This study | Plasmid for the deletion of *BT3091* |
| Plasmid | pLGB13 | pAB007 | This study | Plasmid for the deletion of *BT4581* |
| Plasmid | pLGB13 | pAB008 | This study | Plasmid for the deletion of *BT3087* in Δ*BT3086* |
| Plasmid | pET-29b | pAB010 | This study | Plasmid for the recombinant expression of *BT3086* |
| Plasmid | pET-29b | pAB011 | This study | Plasmid for the recombinant expression of BT3087 |
| Plasmid | pET-29b | pAB012 | This study | Plasmid for the recombinant expression of BT4581 |

**Table 2.**
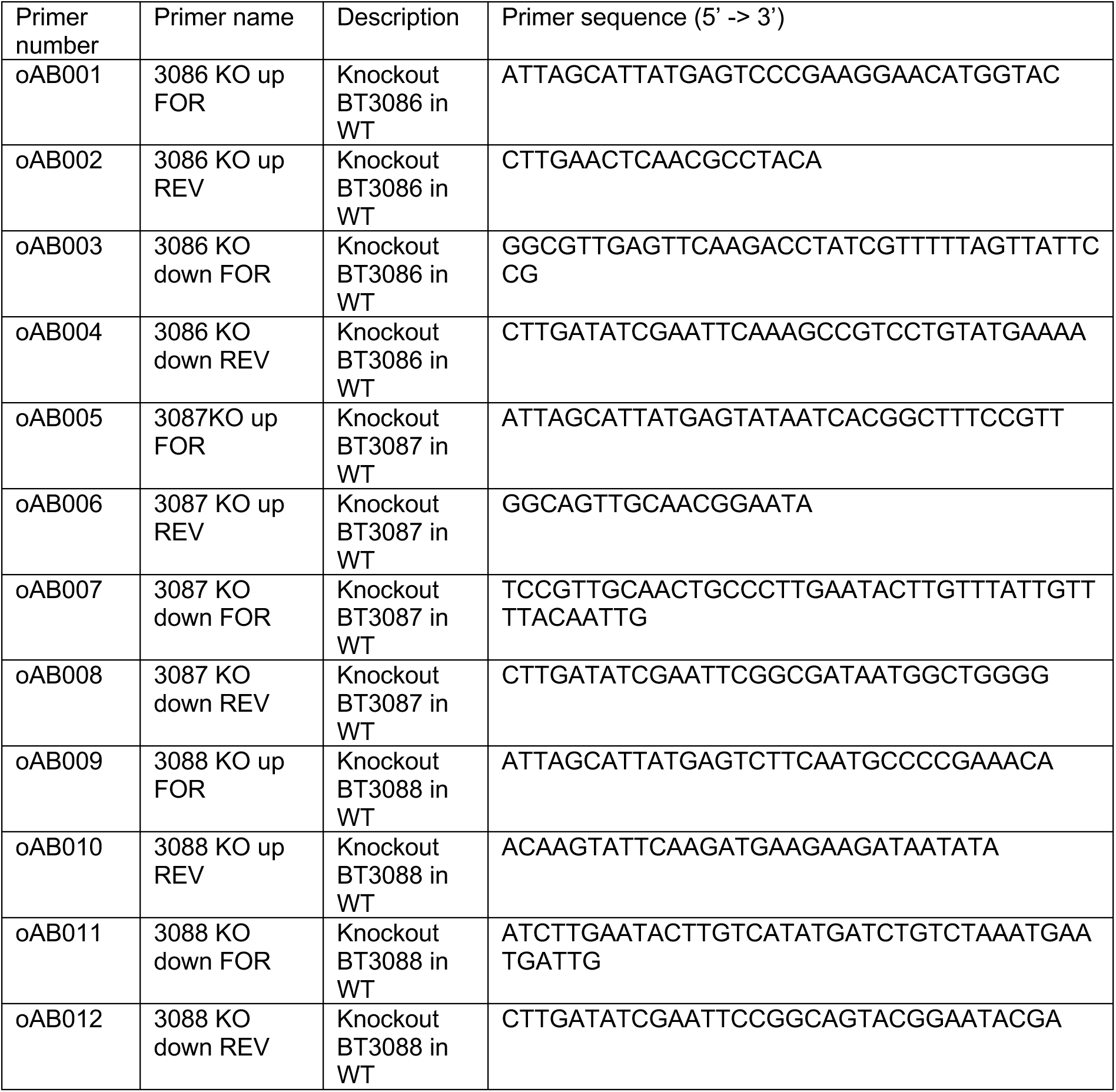
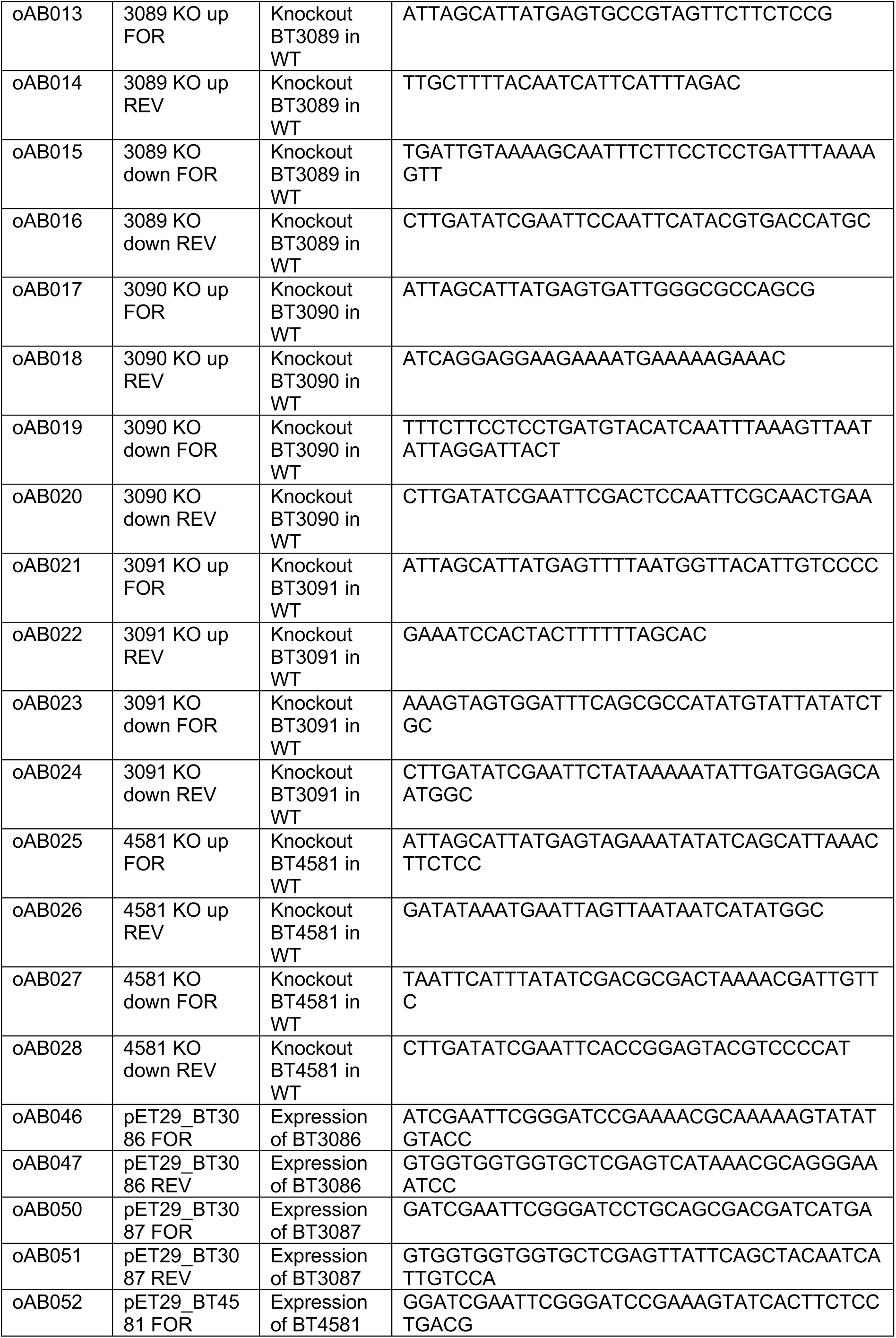
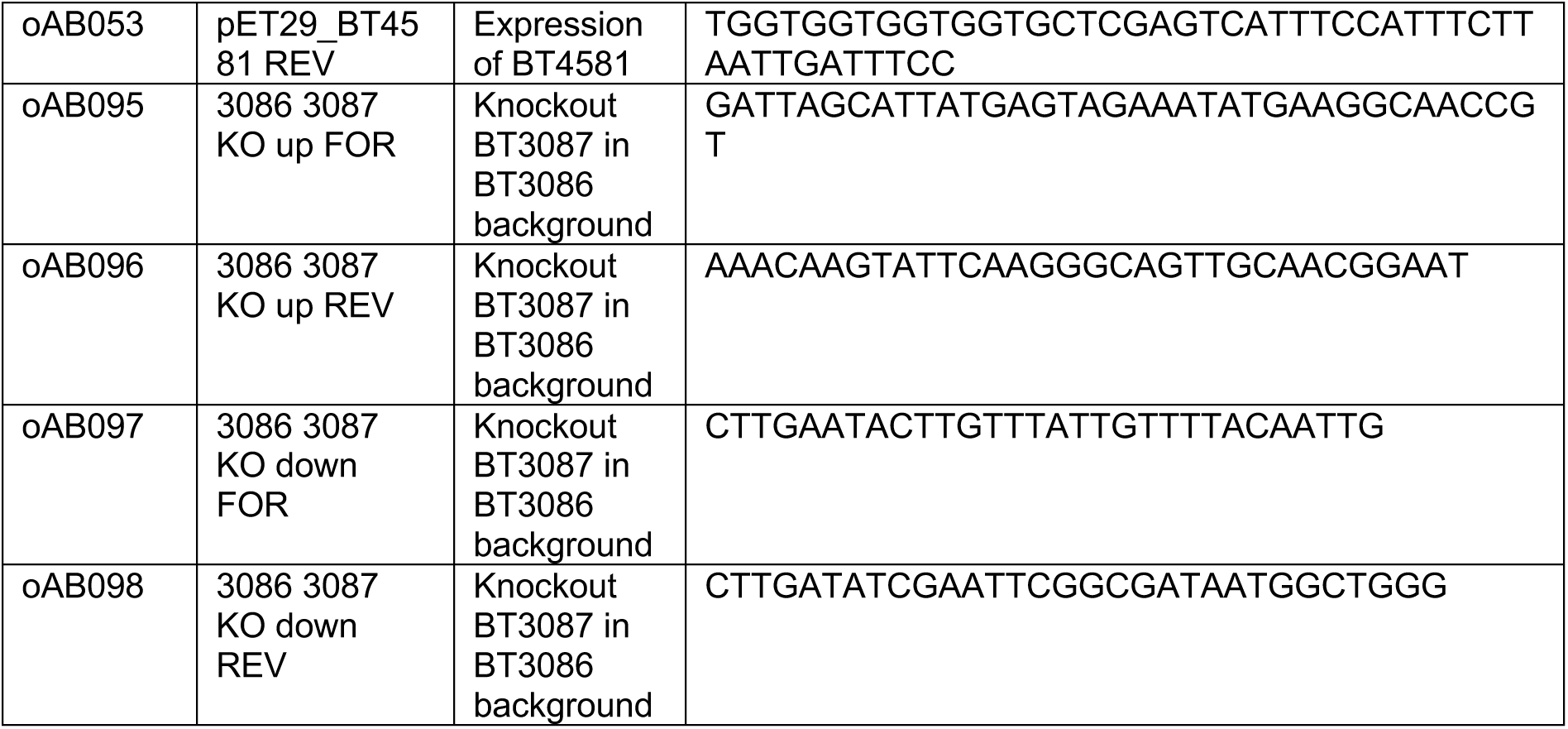
List of primers used for this study.

Given the functional results of the mutants on growth phenotypes, we wanted to further understand the contributions of each individual GHs: BT3086, BT3087 and BT4581 in dextran breakdown. We took an *in-vitro* approach to address this question, where we analysed the degradation products of purified GHs. We first produced the three GHs recombinantly: BT3086 (α-glucosidase II), BT3087 (dextranase) and BT4581 (α-glucosidase), all lacking their signal peptide domains. Individually expressed GH constructs were purified by affinity chromatography and subsequently analyzed by SDS-PAGE (Fig. S1B). We then incubated individual GHs in BMM-dextran. We analyzed the resulting digested carbohydrate contents by HPAEC-PAD. HPAEC-PAD chromatograms of BMM dextran treated by each of the three enzymes all showed similar patterns degradations but with quantitative differences (Fig. 1C). BT4581 produced glucose as the sole degradation product, whereas BT3086 produced glucose and isomaltose as major degradation products. Finally, BT3087 treatment of dextran not only produced glucose and isomaltose as major degradation products, but also other higher MW oligosaccharides (Fig. 1C, D). These data suggest that the enzyme BT3087 is the key GH for the initial debranching of dextran polymers, whereas the enzymes BT3086 and BT4581 plays import roles in liberating glucose from dextran – oligosaccharides, which may be subsequently metabolized by *Bt*.

### Bt growth dynamics depends on dextran MW

Dextran polysaccharides exist in a wide range of MW and branching which influence its complexity ^34,35^. We wondered whether the size of the dextran polymer itself may have an influence on the concerted activity of the sus-like enzymes, and therefore on dextran metabolism and subsequent growth dynamics of *Bt*. To explore this possibility, we grew WT *Bt* in dextrans of 3 different MWs: MW = 9 kDa, 35 kDa and 200 kDa (total mass of dextran is equal between conditions). We cultured *Bt* for 3 days while measuring optical density to eventually generate growth curves (Fig. 2A). We then quantified maximum OD, lag time and growth rates of individual cultures (Fig. 2B). We found that the total biomass *Bt* biomass at its maximum value was independent of dextran MW. However, we found important differences in the dynamics of its growth. First, higher dextran MW tends to increase lag time, thereby delaying recovery from stationary phase. In addition, higher the growth rate of *Bt* is slower in in higher MW dextran. Overall, this leads to a delayed growth pattern for *Bt* in higher MW dextran polysaccharide (Fig. 2A).

**Figure 2:**
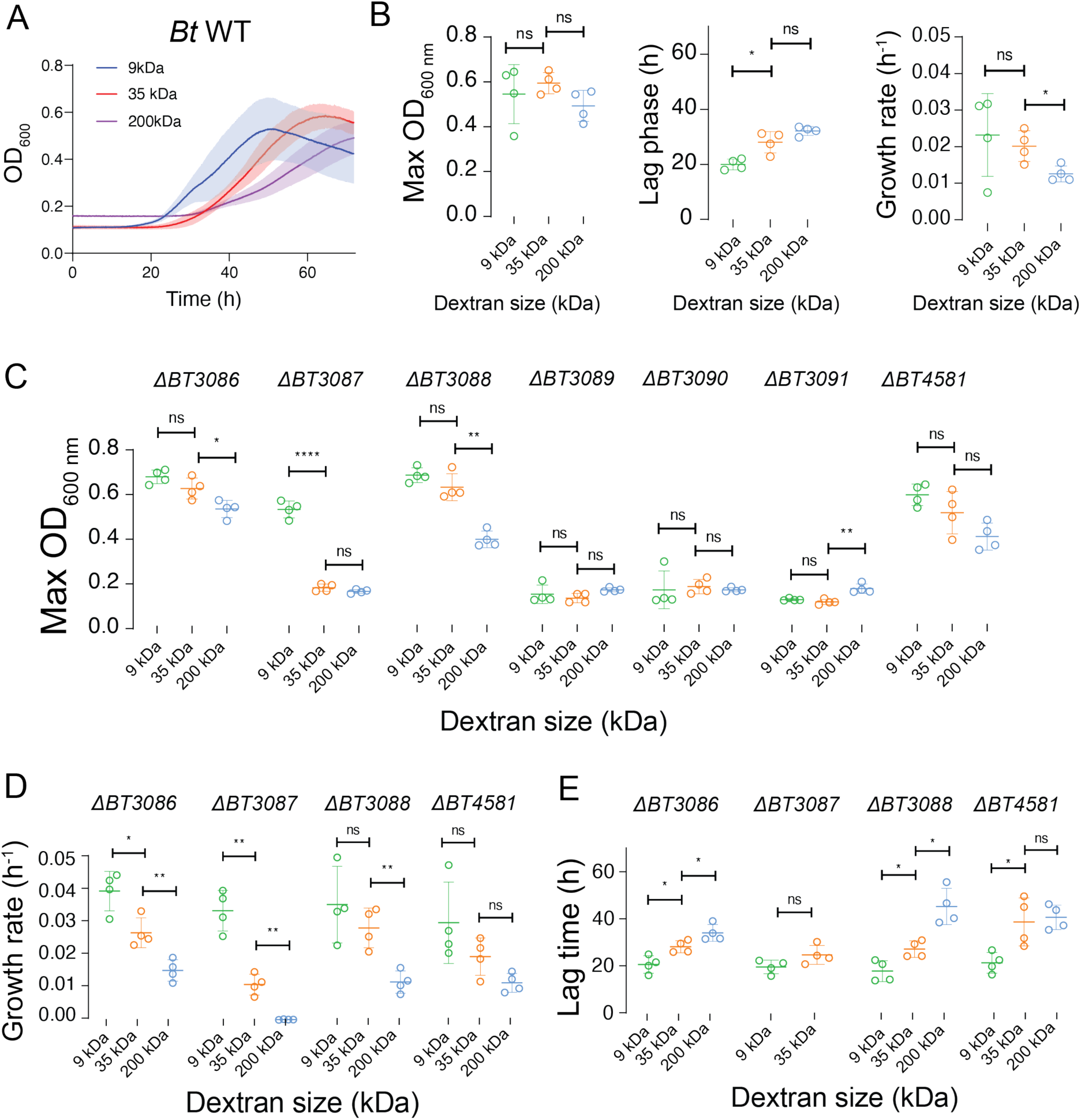
Growth profile quantifications from growth curves of various *Bt* mutants in 4 different sizes of dextran. (A) Growth curves of *Bt* WT in 3 different sizes of dextran. *N = 4.* (B) Extracted parameters from wildtype *Bt* growth curves in dextran. We quantified the maximum OD, lag phases and growth rates of *Bt* WT in 3 different sizes of dextran. We observed a general trend where increasing dextran sizes lead to an increase in lag phase and a decrease in growth rate. (C) Maximum OD of each single mutant cultures in 3 different sizes of dextrans. (D) Quantified growth rates of individual mutants in 3 different sizes of dextrans. Mutants which failed to grow were omitted. (E) Quantified lag time of individual mutant cultures in 3 different sizes of dextrans. Mutants which failed to grow were omitted. (B-E) All growth experiments: *N = 4,* error bars: standard deviation. Statistical test: *t-*tests were performed between pairs of dextran sizes. ns *p* > 0.05, * *p* < 0.05, ** 0.05 < *p* < 0.01, *** 0.001 < *p* < 0.0001 and **** *p* < 0.0001.

As we have already cultured all PUL48 mutants supplemented with dextran 35 kDa in test tubes (Fig. 1B) and observed growth defects in mutants lacking BT3087, BT3089 or BT3090, we then wondered whether changes in dextran MW can further impact the growth dynamics of these *Bt* mutants. To explore this question, we measured the growth of PUL48 gene knockouts supplemented with 3 different MW of dextran polymers with a plate reader (Fig. S2). We first quantified and compared the maximum OD of individual mutant cultures in each of the dextran media (Fig. 2C, S3A). Mutants lacking the GH BT3087 which could not grow in 35 kDa dextran (Fig. 1B), did grow well in 10 kDa dextran. By contrast, we observed that *Bt* mutants lacking BT3089 (SusD), BT3090 (SusC) or BT3091 (SusR) were unable to grow in dextran of any molecular weight. This result suggests that the activity of BT3087 becomes crucial in *Bt* metabolism for growth in high molecular weight dextran. Finally, we observed that the mutant lacking BT3088 (SusE) also grew in all dextran sizes but with weaker growth at 200 kDa, suggesting that this protein also contribute to dextran utilization at high MW.

We wondered whether these changes in biomass were due to perturbations in growth rates or lag time. We thus measured the impact of dextran MW on the growth rate of the various *Bt* mutants by computing the growth rates of all mutants in different sizes of dextran (Fig. 2D, S3C). Like the case of *Bt* WT, we observed a trend of decreasing *Bt* growth rates as dextran MW increased. The mutants lacking BT3087 all led to a decrease in growth rate as dextran MW increased, confirming that BT3087 is a crucial GH for supporting *Bt* growth in high MW dextrans (Fig. S3C). The mutant *ΔBT3088* also experienced a sharp decrease in growth rate between growths in dextran 35 kDa and 200 kDa (Fig. 2D), suggesting the importance of this enzyme in the metabolism of high MW dextran polymers by *Bt*. Finally, we quantified the lag phases of all mutants in different MW of dextran. We observed a general increase in lag time as the size of dextran increases as in WT *Bt.* (Fig. 2E, S3B). The mutant *ΔBT3088* showed the most drastic increase in lag phase between dextran 35 kDa and 200 kDa.

Overall, our data suggest a dextran size dependent growth pattern in *Bt*. As dextran MW increases, this leads to a decreased growth rate of *Bt*. We identified that three enzymes within PUL48: BT3089 (SusD), BT3090 (SusC) and BT3091 (SusR) which are crucial for *Bt* growth in any dextran size. We also found that BT3087 and BT3088 (SusE) promote *Bt* dextran metabolism at high MW. Interestingly, our data also suggest that large dextran oligosaccharides of 9 kDa can be metabolized by *Bt* without the aid of BT3087.

### Tracking dextran binding using fluorescent polymers

To further investigate the mechanisms of binding and import of polysaccharides of various sizes the, we designed a fluorescence microscopy-based assay for dextran localization at the surface of *Bt* cells (Fig. 3A). We cultured *Bt* cells all expressing the fluorescent protein mCherry constitutively in the following backgrounds: WT, *ΔBT3088 (susE-), ΔBT3089 (susD-)* and *ΔBT3090 (susC-).* We grew these strains in BMM containing equal amounts of dextran and glucose. We then diluted the cells in PBS supplemented with fluorescein-conjugated dextran of various sizes. The solution was then incubated anaerobically for one hour, washed and imaged by high-resolution fluorescence microscopy. All samples were imaged in both the red and green fluorescence channels, which correspond to *Bt* cells and fluorescent dextrans respectively (Fig. 3B, S4). Microscopy images of *Bt* incubated with 40 kDa dextran-fluorescein showed a strong localization of green fluorescence signal to WT and *ΔBT3088* cells, but not *ΔBT3089* and *ΔBT3090* cells. These results suggest that mutants lacking SusD or SusC lose the ability to bind 40 kDa dextran on the cell surface, potentially explaining their inability to grow in dextran (Fig. 2C).

**Figure 3:**
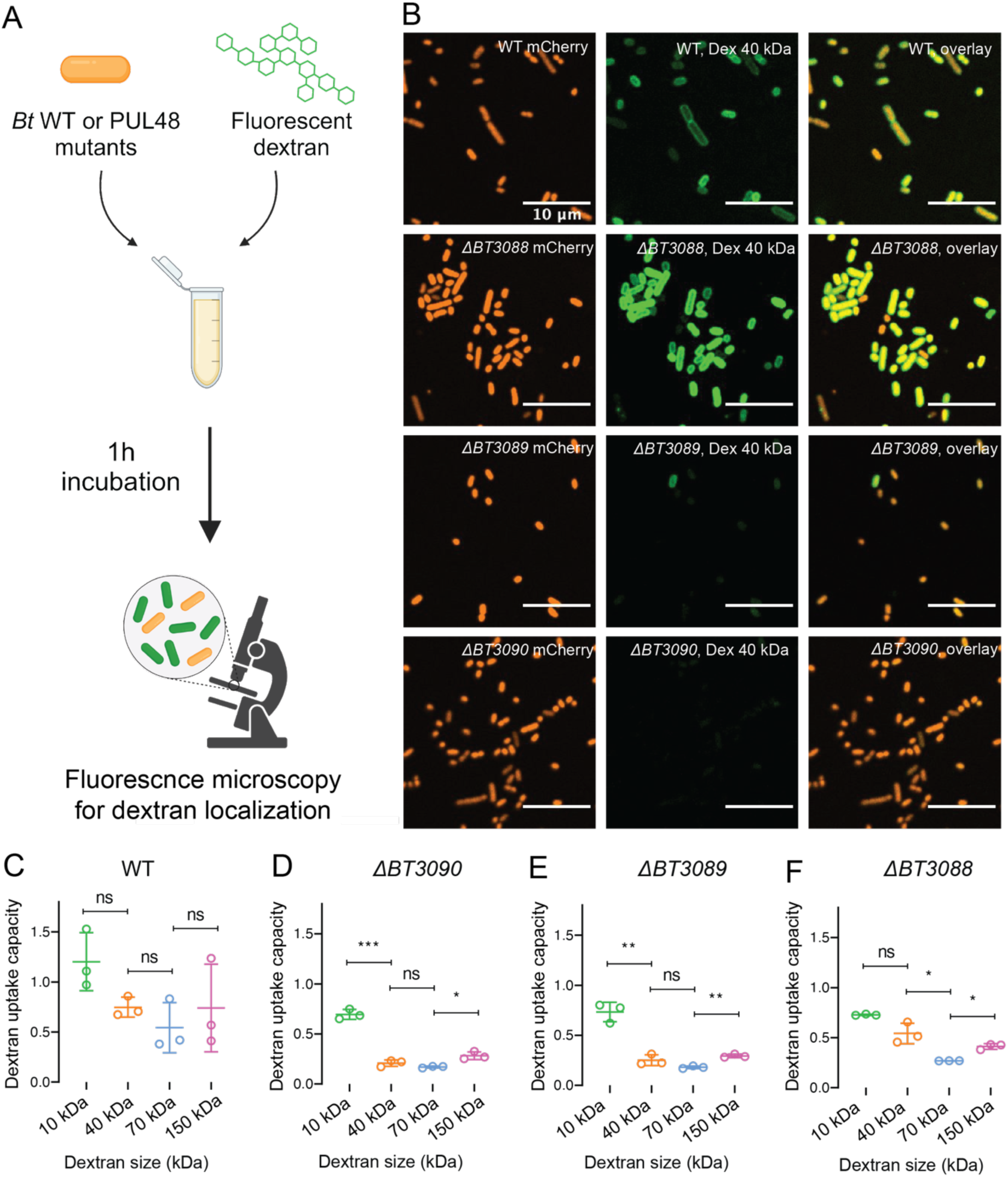
Fluorescent dextran localization to *Bt* cells. (A) Schematic of experimental set-up. *Bt* cells constitutively expressing mCherry are incubated with fluorescein-dextran of various molecular weights for 1 hour, washed and imaged by fluorescence microscopy. (B) fluorescence microscopy images of *Bt* wildtype along with various mutants constitutively expressing mChery incubated in dextran 40 kDa conjugated to fluorescein.Red channel corresponds to *Bt* cells, green channel corresponds to dextran, and the overlay corresponds to localization of dextran to *Bt* cells. (C) The binding capacity of various *Bt* mutants to varying sizes of dextran molecules. The binding capacity is defined as the ratio between green and red fluorescence signals per individual cells. The ratio for individual cells within a field of view were then averaged and plotted. For all fluorescein dextran binding experiments, *N = 3.* Statistical test: *t-*tests were performed between pairs of dextran sizes. * *p* < 0.05, ** 0.05 < *p* < 0.01, *** 0.001 < *p* < 0.0001 and **** *p* < 0.0001.

We then measured the dextran-binding capacity of each *Bt* strain by quantifying dextran fluorescence per cell normalized to the mCherry fluorescence of the same cell. We then average this for all cells. In the case of *Bt* WT, we could not distinguish differences in the dextran binding capacity across the 4 different sizes of dextran polymers tested (Fig. 3C). We then performed the same analysis on PUL48 mutants (Fig. 3D-F). First, we observed that the deletion of BT3089 (SusD) or BT3090 (SusC) led to a significant decrease in green fluorescence upon treatment with dextran polymers of MWs higher than 10 kDa (Fig. 3D, E), suggesting that these two enzymes are important for the specific localization of dextran molecules to the cell membrane of *Bt*. Interestingly, the deletion of BT3088 (SusE) led to a decrease in green fluorescence localization upon treatment with dextran polymers greater than 40 kDa in MW (Fig. 3F), suggesting that this protein is important in *Bt* dextran binding in a size-dependent manner. Finally, we observed that all 3 deletions led to no significant changes in green fluorescence upon treatment with 10 kDa dextran compared to WT (Fig. S5), suggesting that this dextran binds to *Bt*’s cell membrane without the aid of the PUL48 machinery. Together, these data suggest that the PUL48 enzymes BT3088, BT3089 and BT3090 play crucial role in the dextran binding by *Bt*, and that the MW of dextrans itself does not have an impact on the binding capability by WT *Bt* cells.

### PUL48 and dextran size define the extent of nutrient sharing by Bt

We previously demonstrated that *Bt* can share by-products of dextran metabolism with *Bacteroides fragilis* (*Bf*), ultimately guiding community composition and biofilm spatial organization^30^. We thus wondered whether individual PUL48 proteins may influence the nature of nutrient sharing by *Bt* to other members of a community, including in a MW-dependent manner. To answer this question, we grew *Bt*-*Bf* each constitutively expressing distinct fluorescent proteins in competition and monitored the relative fraction of each species in the community using fluorescence microscopy (Fig. 4A). We first quantified overall biomass (via optical density) of the co-cultures after 3 days (Fig. 4B). The growth patterns in co-cultures were identical to the *Bt* monocultures in dextran from Fig. 1C: co-cultures containing mutants which lacks the GH BT3087 or the regulator BT3091 failed to grow. We then measured the abundance of *Bf* in each co-culture by sampling each co-culture upon 1, 2, and 3 days of growth under anaerobic conditions. When computing the *Bf* fraction abundance by computing the ratio between mCherry- and sfGFP-positive cells (Fig. 4C), we found that with when co-cultured with WT *Bt*, the fraction abundance of *Bf* overtime is maintained at 50%^30^. This ratio however decreases in all *Bt* PUL48 mutants which have the ability to grow in dextran: *ΔBT3086, ΔBT3088, ΔBT3089, ΔBT3090, ΔBT4581,* and *Δ86,81*. These results suggest that the deletion of any one of these proteins potentially lead to either a reduction in availability of metabolic by-products like glucose, which ultimately leads to a reduction of nutrient availability for glycolysis in *Bt*.

**Figure 4:**
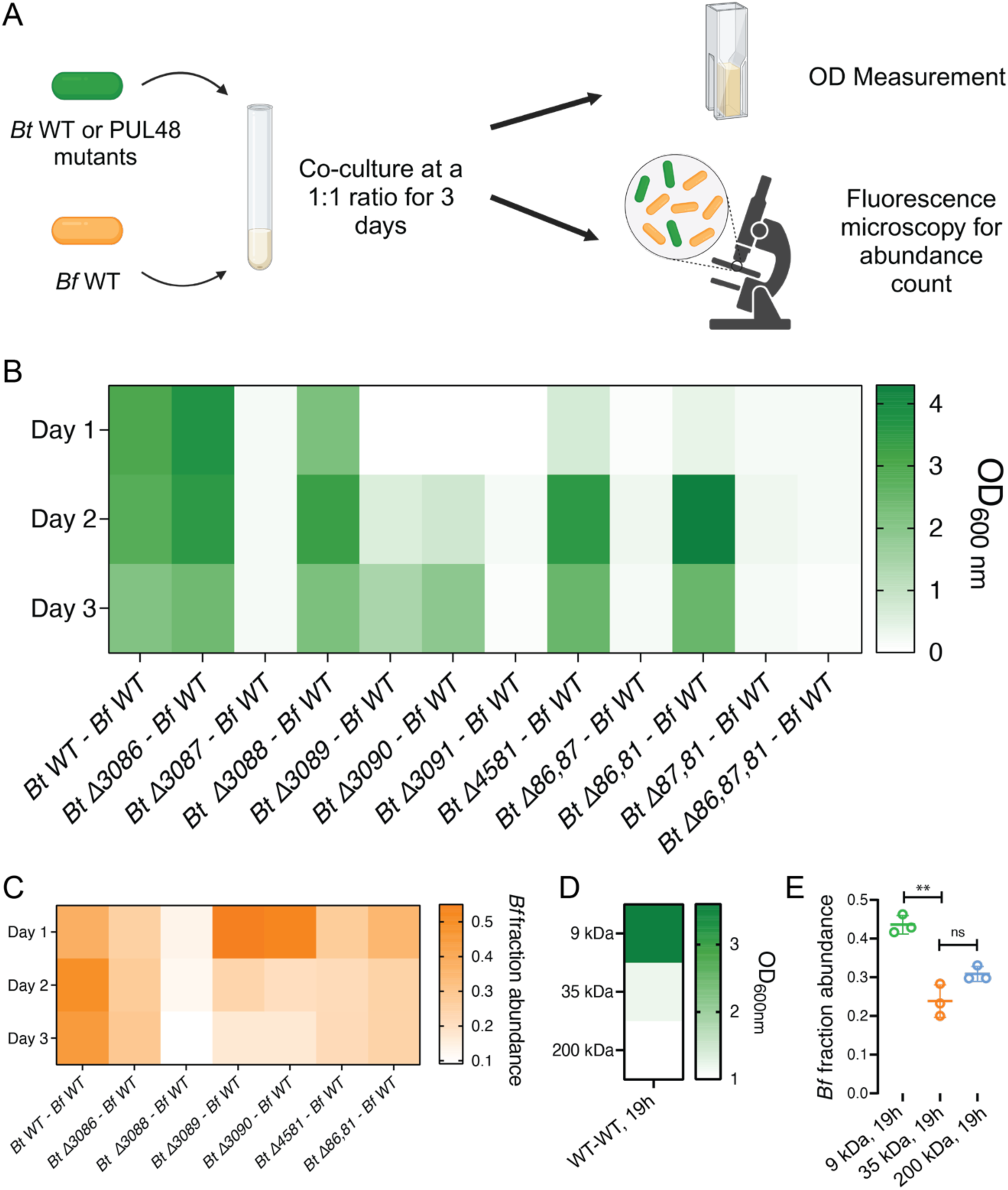
PUL48 proteins contributes to the extent of nutrient sharing from *Bt* to *Bf* in dextran. A) Schematic of experimental set-up. *Bt* WT or mutants expressing sfGFP are mixed with *Bf* WT at a 1:1 ratio. OD measurements and fluorescence microscopy were then performed daily to study coculture growth and *Bf* abundance. (B) Co-culture growths of *Bt* WT or PUL48 mutants with *Bf* in dextran. We monitored the optical density (OD) daily over the course of 3 days. (C) *Bf* fraction abundance from competition experiments. *Bt* strains which failed to grow as monocultures has been omitted from co-culture analysis. Our results indicate that the loss of any one enzyme associated with PUL48 led to a decrease in nutrient sharing to *Bf* in comparison to WT levels. (D) Co-culture growths of *Bt* WT with *Bf* in 3 different sizes of dextran polymers. We monitored the OD of individual cultures upon 19 hours of growth. (E) *Bf* fraction abundance from *Bt* WT - *Bf* WT co-culture experiments in 3 different sizes of dextran upon 19 hours of growth.

Finally, based on different metabolic rates of *Bt*, we hypothesized that the size of polysaccharide polymers modulates nutrient sharing to impact community composition. To address this question, we competed *Bt* WT with *Bf* WT at a 1:1 ratio in 3 different sizes of dextran. We sampled these co-cultures after overnight growth (Fig. 4D), measured optical density along with the fraction abundance of *Bf* by fluorescence microscopy (Fig. 4E). First, we observed that the final OD of co-cultures decrease as the size of dextran polymers increased, consistent with our *Bt* monocultures (Fig. 2A). We found that co-cultures grown in higher MW dextran (35 and 200 kDa) have a decrease in *Bf* relative abundance (Fig. 4E).

These results suggest that the size of polysaccharides impacts *Bt*’s growth rate, where reduced *Bt* growth impacts the amount of glucose metabolites generated at a given time. Increasing dextran MW reduced glucose production rate and secretion which may decrease *Bf* growth. Altogether, these results show that polysaccharide size can impact nutrient sharing in microbial communities, leading to changes in community compositions.

## Discussion

During cycles of dietary shifts, changes in glycan composition greatly contribute to defining microbiota composition^5^. PULs functions are quite specific in processing glycans. However, within each polysaccharide family, these exists a variety of molecular structures introduce another layer of diversity, but how this impacts the functions of PULs has been largely overlooked. To comprehensively understand how diet modulates commensal species metabolism which ultimately shapes our gut microbiota, it is crucial to understand how this physical complexity impacts their metabolic pathways.

We identified a general trend of decreasing *Bt* growth rates as dextran MW increased. We attribute this decrease to the increased structural complexity of dextran polymers through branching, which reduces the processivity of PUL48^36^. As structural complexity increases, the liberation of free oligosaccharides and glucose from dextran may become increasingly difficult for *Bt*. This leads to a reduced pool of oligosaccharides and glucose, leading to an overall decrease in growth rate in dextran polymers of higher MW. At the molecular level, we identified two enzymes whose contribution becomes crucial at high MW: the dextranase BT3087 and the SusE like protein BT3088. We found a cut-off at 9 kDa for BT3087. This differs from the Sus system SusC-SusD complex which selectively imports only short maltoolisaccharides with degree of polymerization of 5 – 16 ^37^, much smaller in comparison to the dextran 9 kDa. This may be attributed to the overall structure of this particular dextran polymer as it is known that dextran 2 to 10 kDa exhibit properties of an expandable coil, with minimal branching^35^, which potentially allow native dextran to pass through the importer BT3090 (SusC) without the aid of BT3087. Another protein which we found to contribute to dextran utilization in a size-dependent manner is BT3088 (SusE). The exact roles of SusE-like proteins polysaccharide utilization have remained unclear^37^.

However, we observed that the deletion of SusE led to a size dependent growth defect in *Bt* when cultured in dextran polymers greater than 35 kDa in size. Our results suggest that BT3088, a SusE like protein, likely plays an important role in *Bt’s* binding to complex dextran polymers or oligosaccharides generated by complex dextran metabolism, which subsequently facilitates the import of extra oligosaccharides into the cell.

Our results also show that dextran surface binding is a key function in its metabolism and depends on surface-exposed GHs. SusC-like and SusD-like proteins plays important roles in binding and uptake. Our fluorescein-dextran binding data for *Δ*BT3088 further confirms that SusE plays an important role in *Bt* binding and accessing dextran greater than 70 kDa in size, suggesting the important role of SusE in the utilization of high molecular weight dextran by *Bt.* Our observations are reminiscent of adhesion of *Ruminococcus bromii*, a common member of the gut microbiota which also utilize starch, which expresses surface enzymes which facilitates adhesion to starch molecules^38^.

We then sort to understand the contribution of polysaccharide molecular weight on species interactions and abundance at the community level. We observed that the deletion of any one protein within PUL48 led to a decrease in *Bf* abundance in a two-species community. Such decrease in *Bf* abundance is likely a result of the reduced availability of the public good glucose. The deletion of BT3086 or BT4581, 2 α-glucosidases which are localized in the periplasm, reduces the formation of glucose from oligosaccharides. As BT3087 also generate glucose as a product, extracellular glucose may be preferentially imported by *Bt*, reducing the overall pool of public goods available for *Bf* to exploit. As the deletion of BT3088 also led to a significant decrease in *Bf* population, BT3088 also likely contribute to the uptake of oligosaccharides in *Bt*, hence also leading to the import of extracellular glucose and reducing nutrient availability to *Bf*.

Previous studied have demonstrated that different types of galactans with differing structural complexities have an impact on the species specific fermentation by gut microbes, which leads to changes in overall microbiota composition^39^. More precisely, it had been previously demonstrated with blackberry polysaccharides that the MW of the polysaccharides impact the rate of fermentation and ultimately the composition of the overall microbial community^40^. Our data also suggests a potential mechanism where polysaccharide size impacts nutrient sharing in microbial communities through defining the growth rate of the utilizer, restricting the amount of public goods generated and shared to the non-utilizer, which ultimately lead to an increasing delay in growth and colonization by the non-utilizer.

Overall, our work provides a novel mechanism of commensal selection based on the complexity of dietary polysaccharides. Our observations imply that at equal molecular composition and concentration, a seemingly identical dietary composition can lead to distinct community composition depending on the structure of polysaccharide molecules it contains. As many other dietary fibres also likely exist in large ranges of molecular weight, understanding how molecular weight impacts utilization by gut microbe may lead to the potential design of next generation precision prebiotics.

## Materials and Methods

### Strains and culture media

*Bacteroides thetaiotaomicron* VPI-5482 WT, *Bacteroides thetaiotaomicron* VPI-5482 sfGFP and *Bacteroides fragilis* NCTC-9343 mCherry, along with all *Bacteroides thetaiotaomicron* VPI-5482 PUL48 knockout mutants (Table 1) were used for experiments in this work. TYG medium and *Bacteroides* minimal medium (BMM) supplemented with the indicated carbon source were used in all experiments.

For TYG medium, 10g tryptone, 5g yeast extract, 2.5g D-glucose and 0.5g L-Cysteine were dissolved in 465 mL of ddH_2_O. The resulting solution was then immediately autoclaved and allowed to cool down to room temperature. In the meantime, Salt Solution A was prepared by mixing 0.26g of CaCl_2_ and 0.48g of MgSO_4_ in 300 mL of ddH_2_O until fully dissolved where 500 mL of ddH_2_O was then added. 1g KH_2_PO, 1g K_2_HPO_4_ and 2g of NaCl were then added to the solution and stirred at room temperature until all salts are fully dissolved followed by the addition of 200 mL of ddH_2_O to top up the volume. The salt solution was then stored at 4°C for future use. A 10% (w/v) NaHCO_3_ solution was prepared by dissolve 50g of NaHCO_3_ into 500 mL of ddH_2_O followed by filter sterilization with a 0.22 µm filter. A 1.9 mM Hematin – 0.2M Histidine solution was prepared by dissolving 60.18 mg of hematin in 1 mL of 1M NaOH until fully solubilized, then neutralized with 1 mL of 1M HCl. In a separate beaker, 1.55 g of L-Histidine was dissolved in 48 mL of ddH_2_O. The histidine solution was then combined with the hematin solution, filter sterilized with a 0.22 µm filter and stored at 4°C for future usage. 5 mL of hematin-histidine solution and 10 mL of 10% NaHCO_3_ solution were added to the TYG base medium with a 0.22 µm filter. With a separate 0.22 µm filter, 20 mL of Salt Solution A was then added. The resulting complete TYG medium was then stored at 4°C for future usage.

The BMM medium was prepared as previously described^41,42^. Briefly, a 10x medium stock containing 1 M KH_2_PO_4_, 150 mM NaCl and 85 mM (NH_4_)_2_SO_4_ was first prepared and adjusted to a pH of 7.2. Solutions of 1 mg/mL vitamin K_3_, 0.4 mg/mL FeSO_4_, 0.1 M MgCl_2_, 0.8% (w/v) CaCl_2_, 0.01 mg/mL vitamin B_12_ and 1.9 mM hematin–0.2 M histidine solutions were prepared separately. To prepare 100 mL of BMM,10 mL of the 10x salts were mixed with 0.1 g of L-cysteine, 50 μL of vitamin B_12_ solution, and 100 μL of each vitamin K_3_, FeSO_4_, MgCl_2_, CaCl_2_ and hematin-histidine solution. All carbon sources were added to a final concentration of 5 mg/mL. The media were filter-sterilized using a 0.22 μm filter unit and degassed in the anaerobic chamber overnight.

All bacteria were cultured at 37 °C under anaerobic conditions in a vinyl anaerobic chamber (COY) inflated with a gas mix of approximately 5% carbon dioxide, 90% nitrogen and 5% hydrogen (CarbaGas). Bacteria were first streaked on brain heart infusion plates (SigmaAldrich) containing 10% sheep blood (TCS BioSciences). Single colonies were then inoculated in BMM and grown overnight.

### Recombinant expression and purification of Bt GHs

The amino acid sequences of the GHs BT3086, BT3087 and BT4581 were collected from the *B. thetaiotaomicron* VPI-5482 genome. The amino acid sequences of each GHs were first analyzed by SignalP^43^ in order to identify signal peptide and potential transmembrane domains of the GHs. Each of these genes were then amplified by PCR without the corresponding signal peptide regions and cloned into the expression vector pET29b containing an N-terminal 10x His-tag. The vector was then transformed into *E. coli* XL10 gold for plasmid production and storage. The resulting plasmid was then transformed into *E. coli* BL-21, where then a 50 mL overnight culture is grown in LB-kanamycin (Kan). The overnight culture was then transferred into 2 L of LB-kan, and grown until the OD of 0.6 at 37 °C. The cells were then induced with 1M IPTG and cultured overnight at 19 °C. The culture was then pelleted, washed, lysed and centrifuged at 20000 rpm to remove membrane material. The resulting supernatant was then purified using an affinity Ni-NTA column, eluted in 1x PBS, and stored at −80 °C until usage. The resulting purified protein were then analyzed by SDS-PAGE to confirm expression and identity.

### GHs activity analysis by HPAEC-PAD

20 µL of 1 mg/mL purified GHs: BT3086, BT3087 or BT4581 were added to 1 mL of BMM-dextran M_w_ 9000-11000. The mixture was then incubated overnight anaerobically at 37°C. Cation and anion exchange chromatography with AmberChrom resins (SigmaAldrich) were then used to remove protein and ions from the medium, followed by filtering through a 0.22 µm sterilization filter prior to analysis. The resulting flow-throughs were then analyzed by HPAEC-PAD. The sugars were separated on a Dionex CarboPac PA-20 column from Thermo Fisher Scientific with the following eluent: A) 100 mm NaOH; B) 150 mm NaOH and 500 mm sodium acetate at a flow rate of 0.4 mL/min. The gradient was 0 to 15 min, 100% A (monosaccharide elution); 15 to 26 min, gradient to 10% A and 90% B (malto-oligosaccharide elution); 26 to 36 min, 10% A, 90% B (column wash step); and 36 to 46 min step to 100% A (column re-equilibration). Peaks were identified by co-elution with known glucose, isomaltose standards using the Chromeleon software. The HPLC data was then exported and replotted with the graphing software GraphPad Prism 8, where area under the curve of peaks of corresponding samples were calculated.

### In frame deletion knockouts in Bt

*Bt* knockouts were generated as previously described^44^. Briefly, 700 bp upstream and downstream of each gene of interest were amplified from *Bt* WT by PCR. The resulting PCR products were then analyzed on a 1% agarose gel and subsequently gel-extracted. The knockout plasmid pLGB13 was then restriction digested with PstI and BamHI, where the upstream fraction, downstream fraction and digested pLGB13 were assembled via Gibson Assembly. The assembled plasmid was then transformed into the donor strain *E. coli S17*. The donor strains carrying the knockout plasmids were then grown to an OD of 1-1.2 in LB-Amp 100. At the same time, *Bt* strains were grown anaerobically to an OD of roughly 0.1.

The donor culture was then washed with PBS, pelleted, and mixed with the *Bt* culture. The culture is then puddled onto BHI plates supplemented with 10% sheep-blood and dried for conjugation aerobically overnight at 37 °C. The resulting carpet is then resuspended in TYG medium and restreaked onto BHI sheep blood plates with gentamycin (Gm) 200 µg/mL and erythromycin (Er) 20 µg/ uL and grown anaerobically for 30 hours. Resulting colonies were picked and restreaked directly onto another BHI-sheep blood plate containing Gm/Er for another 30 hours anaerobically. Resulting single colonies were then picked and restreaked onto BHI-sheep blood plate containing 0.1 µg/mL of anhydrotetracycline (aTc) for counter-selection and grown anaerobically for 30 hours. Resulting single colonies were again picked, where colony PCR is then performed, and the resulting PCR product was sequenced to confirm whether the knockout was successful.

### Insertion of sfGFP and mCherry into Bt and Bf

Fluorescent strains of *Bt* and *Bf* were generated as described previously.^45^ Briefly, the plasmids pWW3452 (GFP) and pWW3515 (mCherry) were transformed into the donor strain *E. coli S17*. The donor was then grown to an OD of 1-1.2 in LB-Amp 100. At the same time, *Bt* or *Bf* strains were grown anaerobically to an OD of roughly 0.1 (for *Bf,* OD 0.01). The donor culture was then washed with PBS, pelleted, and mixed with either the *Bt* or *Bf* culture. The culture is then puddled onto BHI plates supplemented with 10% sheep-blood and dried for conjugation aerobically overnight at 37 °C. The resulting carpet is then resuspended in TYG medium and restreaked onto BHI sheep blood plates with gentamycin (Gm) 200 ug/mL and erythromycin (Er) 20 ug/ uL and grown anaerobically for 30 hours. Resulting colonies were picked and restreaked directly onto another BHI-sheep blood plate containing Gm/Er for another 30 hours anaerobically. Resulting colonies were then resuspended and screened for fluorescent properties on the confocal fluorescence microscope.

### Test tube mutant growths and competitions

For initial growth experiments, *Bt* and various mutants were cultured in BMM supplied with 5 mg/mL of D(+)-glucose (Carl Roth) overnight anaerobically. The resulting cultures were then pelleted by centrifugation and washed twice with PBS. The pellets were then resuspended in PBS, where the OD was measured. The various *Bt* strains were then diluted to a final OD of 0.1 or 0.01 in BMM dextran (M_W_ = 35 kDa). The cultures were then grown anaerobically for 3 days, where the OD of individual mutants were measured after day 1, 2 and 3. For *Bt* mutant competition experiments with *Bf*, *Bt* and *Bf* were cultured separately in 1 mL of BMM-glucose overnight. The resulting cultures were then pelleted by centrifugation and washed twice with PBS. The pellets were then resuspended in PBS, where the OD was measured. *Bt* and *Bf* were then mixed at a 1:1 ratio to an initial OD of 0.1 in each of the carbon sources tested. The cultures were then grown anaerobically for 3 days. A microscopy approach was then used to quantify the relative abundances of *Bt* and *Bf* in these cultures. To achieve this, the cultures were mixed well and 10 µL of each cell culture was sampled at day 1, 2 and 3. The cultures were then diluted in PBS and incubated at aerobic conditions. The samples were then spotted onto a glass coverslip and covered by a thin agarose pad prior to fluorescence imaging by fluorescence microscopy. For each sample and replicate, 3 image frames across the agar pad were recorded. The relative abundance of sfGFP- and mCherry-expressing cells in each sample were then quantified using the ‘analyze particle’ function in Fiji and averaged across the 3 frames.

### Growth curves analysis of Bt

*Bt* and various mutants were cultured in BMM supplied with 5 mg/mL of D(+)- glucose overnight anaerobically. The resulting cultures were then pelleted by centrifugation and washed twice with PBS. The pellets were then resuspended in PBS, where the OD was measured. The various *Bt* strains were then diluted to a final OD of 0.1 in BMM dextran of the corresponding molecular weights in a clear COSTAR 96 wells plate with a final volume of medium at 200 µL. The plate is then completely sealed with parafilm, moved into the Tecan Safire^2^ plate reader, and grown anaerobically at 37 °C. settings: kinetic, 5 min cycles, 875 cycles, shake 10 seconds prior to each analysis cycle. The resulting data were then rearranged by a house-built python script, where all growth curves were then plotted by the graphing software GraphPad Prism 8. For further analysis for the lag time, max OD and maximum growth rate of each conditions, the file of rearranged data was analyzed using Pyphe growth curves^46^.

### Bt binding to fluorescently-labelled dextran

*Bt* WT and various mutants containing mCherry were cultured in BMM supplied with 2.5 mg/ mL of D(+)-glucose and 2.5 mg/mL of dextran 35 kDa overnight anaerobically. The resulting cultures were then pelleted by centrifugation and washed twice with PBS. The pellets were then resuspended in 1x PBS, where the OD was measured. We then diluted the cells to a final OD of 0.1 in PBS and added fluorescein-conjugated dextran of various sizes (Sigma-Aldrich) to the solution at a final concentration of 1 mg/mL. We then incubated the solution anaerobically for 1 hour, and the cells were then washed 3x in 1x PBS. Resulting cells were then resuspended in 1x PBS and imaged by fluorescence confocal microscopy. For each sample and replicate, 5 image frames across the agar pad were recorded. The relative abundance of mCherry-expressing cells sfGFP signal corresponding to dextran binding and import were recorded for each single cell within each samples using the ‘analyze particle’ function in Fiji. The ratio of sfGFP:mCherry were then quantified for every individual cell within an experiment and averaged per experiment.

### Microscopy

All imaging were performed using a Nikon Eclipse Ti2-E inverted microscope coupled with a Yokogawa CSU W2 confocal spinning disk unit and equipped with a Prime 95B sCMOS camera (Photometrics). The 40 x Plan APO objective with a numeric aperture (N.A.) of 0.9 was used for all imaging. Fiji was then used to display the images.

## Data availability

The macro used for analyzing green fluorescence localization to red fluorescing cells will be available on GitHub (https://github.com/PersatLab/Metabolism).

## Acknowledgements

We thank Kelvin Lau at the protein production and structure core facility at the EPFL for help with tips for GH expression constructs, Leonor Garcia-Bayona for tips with *Bt* knockouts, and Lorenzo Talà for help with image analysis. This work was supported by SNSF Projects grant 310030_204190, the Human Frontier Science Program grant number RGY0077/2020 and the EPFL iPhD program.

## Author contributions

J.P.H.W., T.J.B and A.P. conceived the study. J.P.H.W. and A.P. designed the study. J.P.H.W. and N.C. performed all experiments. J.P.H.W., M.F.S. and S.C.Z performed HPAEC-PAD. J.P.H.W and N.C. performed all image and data analysis. J.P.H.W created the figures. J.P.H.W., T.J.B. and A.P. supervised the work. J.P.H.W. and A.P. wrote the paper. All authors provided input on the paper.

---

**Figure S1:**
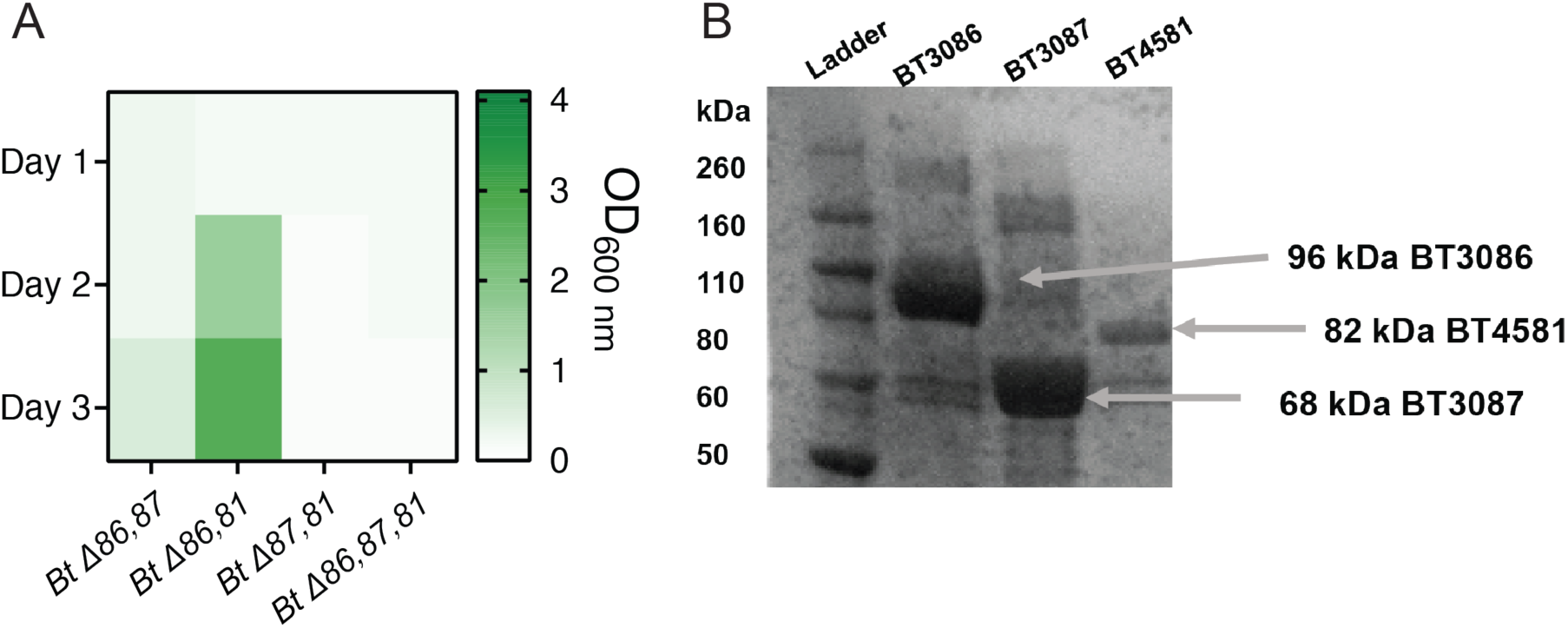
Involvement of GHs BT3086, BT3087 and BT4581 in *Bt* dextran metabolism. (A) Monoculture growths of *Bt* double and triple GH knockout mutants. We measured the optical density (OD) of individual cultures everyday over the course of 3 days (B) SDS-PAGE analysis of recombinantly expressed BT3086. BT3087 and BT4581 upon purification by Ni-NTA affinity chromatography.

**Figure S2:**
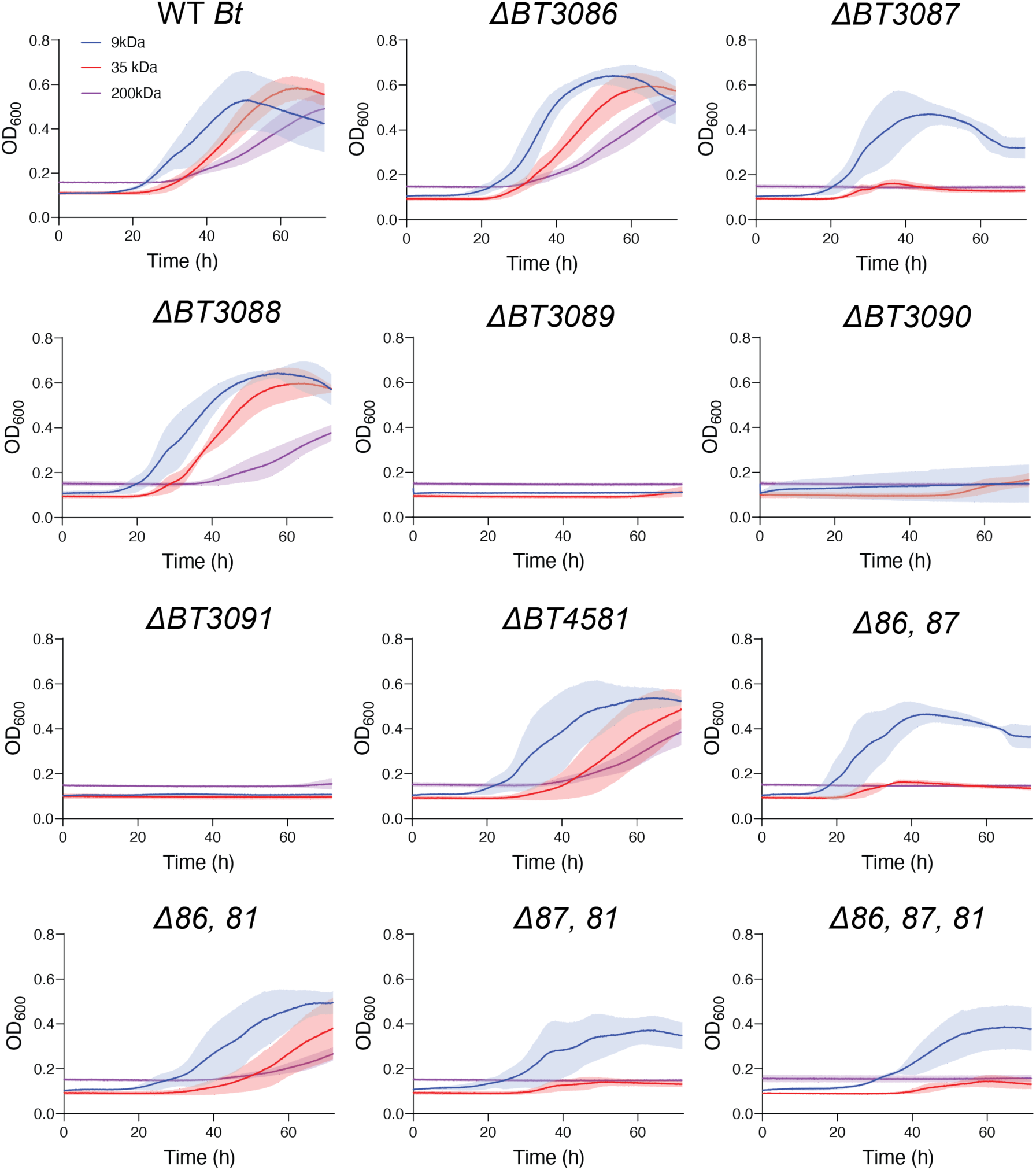
Growth curves of *Bt* WT and various PUL48 mutants in 3 different sizes of dextran over the course of 3 days. *N = 4,* shaded region: standard deviation.

**Figure S3:**
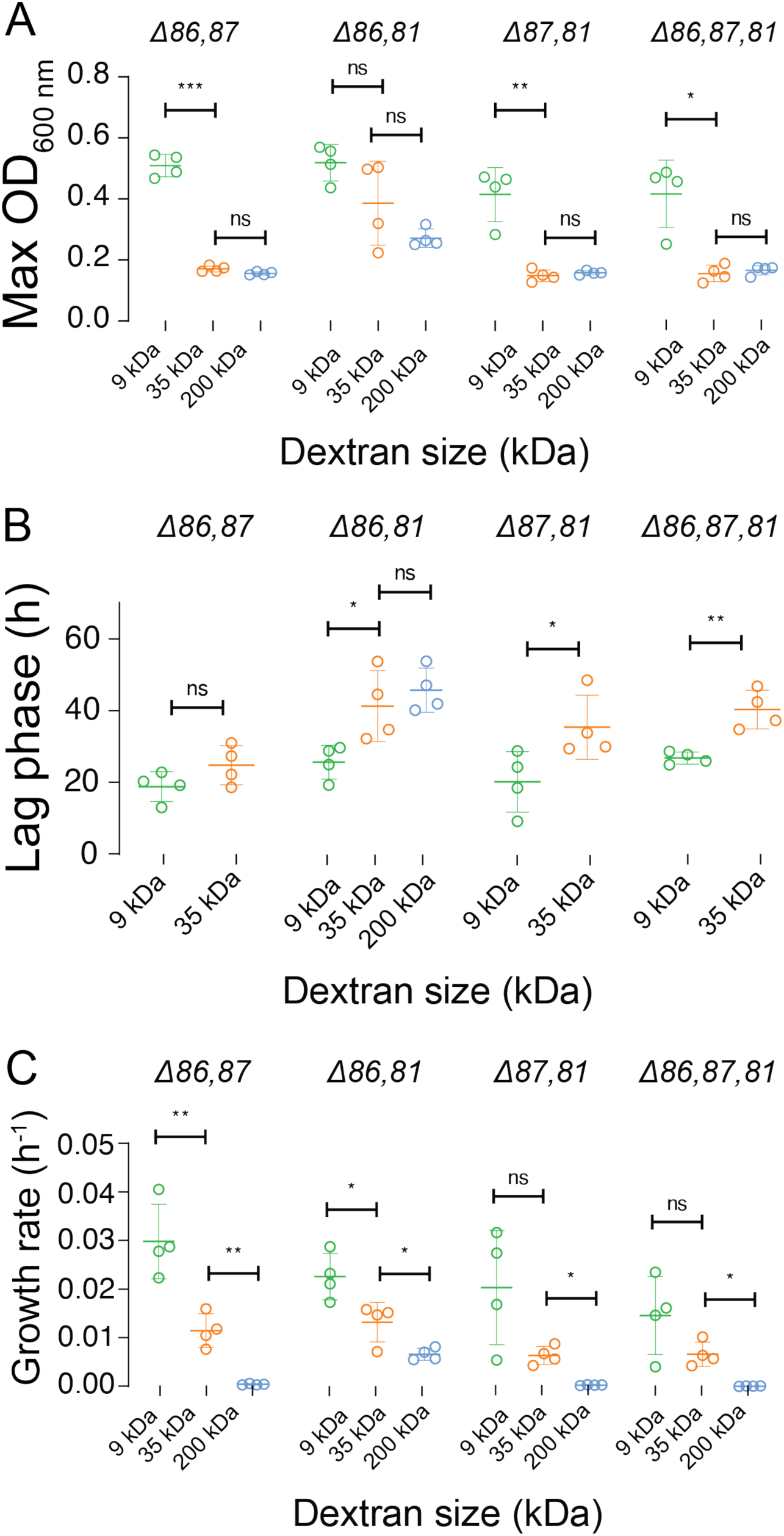
Growth of *Bt* double and triple GH mutants in 3 different sizes of dextran. (A) Quantified maximum OD of each mutant monocultures in dextrans (B) Quantified lag phase of double and triple GH mutants in 3 different sizes of dextrans. (C) Quantified growth rates of double and triple GH mutants in 3 different sizes of dextrans. All growth experiments: *N = 4,* error bars: standard deviation. Statistical test: *t-*tests were performed between pairs of dextran sizes. ns *p* > 0.05, * *p* < 0.05, ** 0.05 < *p* < 0.01, *** 0.001 < *p* < 0.0001.

**Figure S4:**
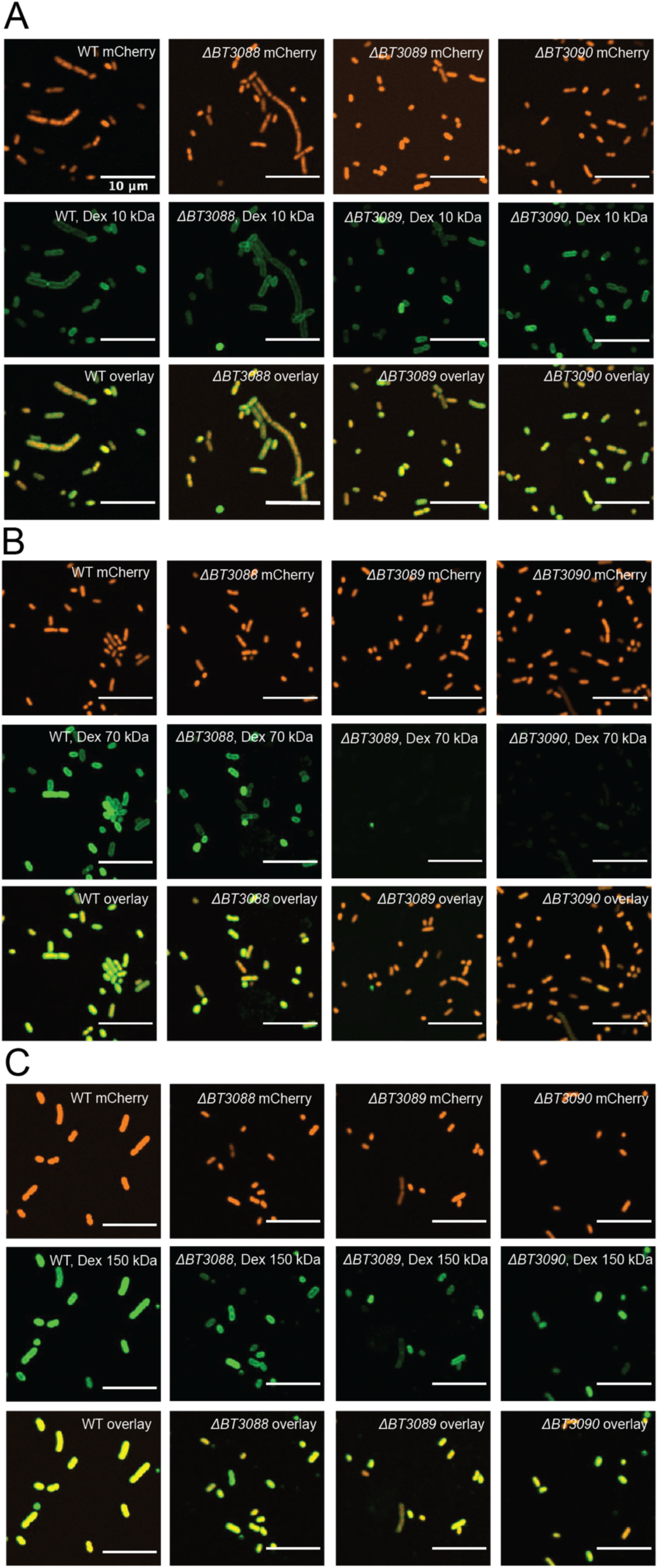
Visualization of dextran localization to *Bt* cells. Fluorescence microscopy images of *Bt* wildtype along with various mutants constitutively expressing mChery incubated in (A) dextran 10 kDa, (B) dextran 70 kDa, and (C) dextran 150 kDa. All dextrans conjugated to fluorescein.

**Figure S5:**
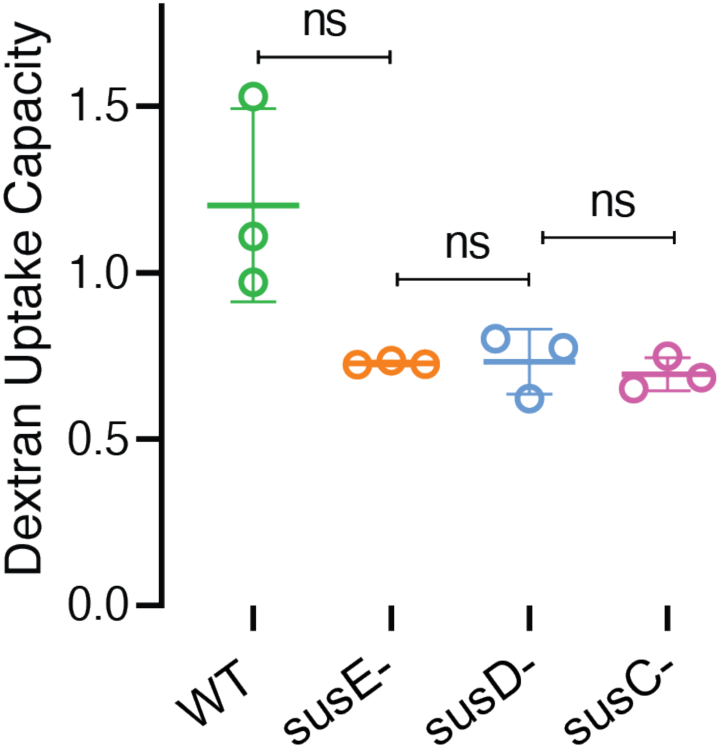
Dextran uptake capacity of 10 kDa dextran by *Bt* WT and 3 PUL48 mutants. No significant difference in the uptake capacity of dextran 10 kDa was observed across the 4 strains analyzed. *N = 3*, Statistical test: Statistical test: unpaired t-tests, ns *p* > 0.05.

